# SNVformer: An Attention-based Deep Neural Network for GWAS Data

**DOI:** 10.1101/2022.07.07.499217

**Authors:** Kieran Elmes, Diana Benavides-Prado, Neşet Özkan Tan, Trung Bao Nguyen, Nicholas Sumpter, Megan Leask, Michael Witbrock, Alex Gavryushkin

## Abstract

Despite being the widely-used gold standard for linking common genetic variations to phenotypes and disease, genome-wide association studies (GWAS) suffer major limitations, partially attributable to the reliance on simple, typically linear, models of genetic effects. More elaborate methods, such as epistasis-aware models, typically struggle with the scale of GWAS data. In this paper, we build on recent advances in neural networks employing Transformer-based architectures to enable such models at a large scale. As a first step towards replacing linear GWAS with a more expressive approximation, we demonstrate prediction of gout, a painful form of inflammatory arthritis arising when monosodium urate crystals form in the joints under high serum urate conditions, from Single Nucleotide Variants (SNVs) using a scalable (long input) variant of the Transformer architecture. Furthermore, we show that sparse SNVs can be efficiently used by these Transformer-based networks without expanding them to a full genome. By appropriately encoding SNVs, we are able to achieve competitive initial performance, with an AUROC of 83% when classifying a balanced test set using genotype and demographic information. Moreover, the confidence with which the network makes its prediction is a good indication of the prediction accuracy. Our results indicate a number of opportunities for extension, enabling full genome-scale data analysis using more complex and accurate genotype-phenotype association models.

## 1. Introduction

Genome-wide association studies over large cohorts are the most powerful approach for identifying genotypes associated with phenotypes. As an example, studying urate and gout, GWAS have identified the major urate transporters and transcription factors involved in the kidney (Leask & Merriman, 2021; Boocock et al., 2020). Additionally, estimated effects of causal variants from GWAS can improve prediction of phenotypic outcomes. Thus the availability of large data sets has brought a new era, with the development of predictive models for complex diseases using genotypes and clinical variables. However, with respect to gout and its related traits, only a small handful of studies have attempted to predict the phenotype using genotypes (genetic risk score) and demographics, and with varying accuracy (Tin et al., 2019; Zhang & Lee, 2020; Sun et al., 2022).

Prediction of a trait is limited by two main factors: heritability of the trait and accurate estimation of the underlying genetic effects (Wei et al., 2014). Thus, current methods underlying the generation of GWAS data (e.g. at the UK Biobank (UKBB) (Sudlow et al., 2015)) bring challenges for prediction, i.e. it is typical to opt for simple but biologically unrealistic models, such as the linear model, in data analyses to enable genome-wide screens for single nucleotide variants (SNVs) with significant effects on (disease) phenotypes. As a result, most GWAS assume the independence of individual SNV effects on the phenotype, and therefore cannot model gene-gene interactions (epistasis). Additionally, typical GWAS are done using a simple additive model, that fails to account for potential dominance effects of different genotypes (Zhu et al., 2015; Hivert et al., 2021). They also often simplify complex multi-dimensional phenotypes to a one-dimensional response variable (Lienkaemper et al., 2018). If epistasis contributes to the genetic architecture of a trait, then identifying epistatic variants is important for improving the predictive power of a GWAS (Crona et al., 2017). Current methods for detecting epistatic interactions are limited computationally and statistically, and developing methods to overcome these challenges is difficult (Wei et al., 2014; Elmes et al., 2021b;a).

Deep neural networks, including Transformer-based ones initially designed for natural language processing (NLP) tasks (Vaswani et al., 2017), have recently been successfully employed to analyse molecular sequence data, the most successful and widely-known example being AlphaFold-2 (Jumper et al., 2021). These approaches have shown great promise in both scalability and potential to support complex models, such as residue-residue interactions in proteins.

Presently, however, such architectures are capable only of handling sequences of modest length, such as those encoding specific proteins. Recently, and in parallel, Transformer-based architectures for NLP have been developed that aim to accommodate extended bodies of text (Wang et al., 2020; Choromanski et al., 2020). Here, we combine these two directions of research and present a Transformer-based deep neural architecture for GWAS data, including a purpose-designed SNV encoder, that is capable of modelling gene-gene interactions and multidimensional phenotypes, and which scales to the whole-genome sequencing data standard for modern GWAS. We trained our model on data from the UK Biobank (UKBB)(Sudlow et al., 2015) to predict gout; that is, we used genotyped SNV data as an input and the patient’s gout diagnosis as an output. The training set was composed of 12,108 cases, of which 6,054 had gout and 6,054 did not.

### 1.1. Transformers background

Neural networks using Transformer blocks have been applied not only to natural language processing tasks (Devlin et al., 2018) but also to the protein folding problem (Jumper et al., 2021) and prediction of nucleotides in DNA sequences based on upstream and downstream context (Ji et al., 2021). The key novelty of the Transformer block is the use of the attention mechanism (Vaswani et al., 2017), where self-attention “heads” assign a relevance score for every token in the sequence with respect to the rest of the tokens via attention calculations. These calculations work by projecting token vectors onto *d*-dimensional key **K**, query **Q**, and value **V** vectors, then taking the following dot products of these projections for each head.

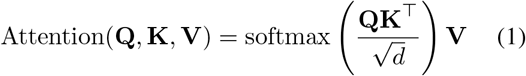

The time and space complexity of attention calculation in Eq. 1 is quadratic in the input sequence length *n*. Significant efforts have recently been made to reduce this cost (Zaheer et al., 2020; Rae et al., 2019; Beltagy et al., 2020). One of the proposed solutions is the Linformer model (Wang et al., 2020), which we build on in this study. As a result of a singular value decomposition analysis on the attention calculation, it is demonstrated (Wang et al., 2020) both theoretically and experimentally that the attention calculation can be reduced from quadratic to linear complexity by including two learnable low-dimensional projections over Eq. (1).

## 2. Methodology

### Data

We obtained SNV data for approximately 487, 000 individuals from the UKBB data bank. For each individual, around 870, 000 SNVs were measured, and a further 90 million imputed. We pre-process data in three steps: 1) extract a sequence of SNVs, with corresponding genome positions, for each person, 2) build a mapping from each SNV to an integer as per Table 1, 3) use this mapping to construct a sequence of integers representing the SNV sequence for each person – see the top of Figure 1.

**Table 1:**
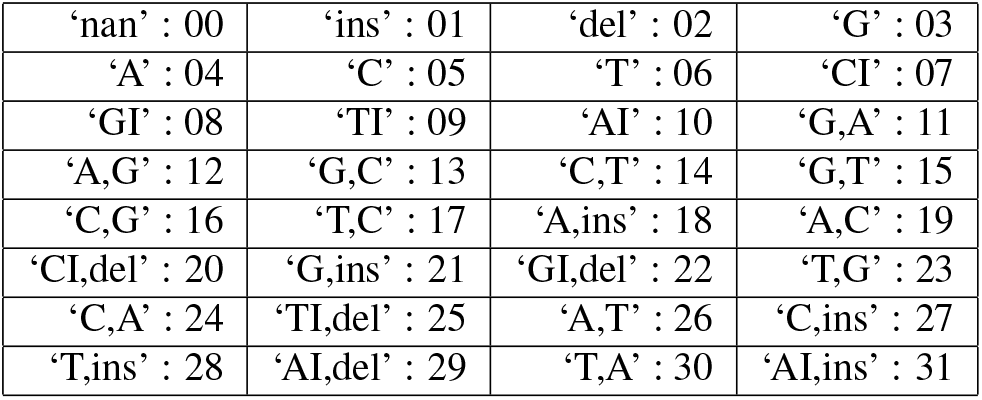
SNV encoding. Homozygous alleles are encoded as ‘X’, heterozygous as ‘X,Y’, I encodes unique sequences.

**Figure 1:**
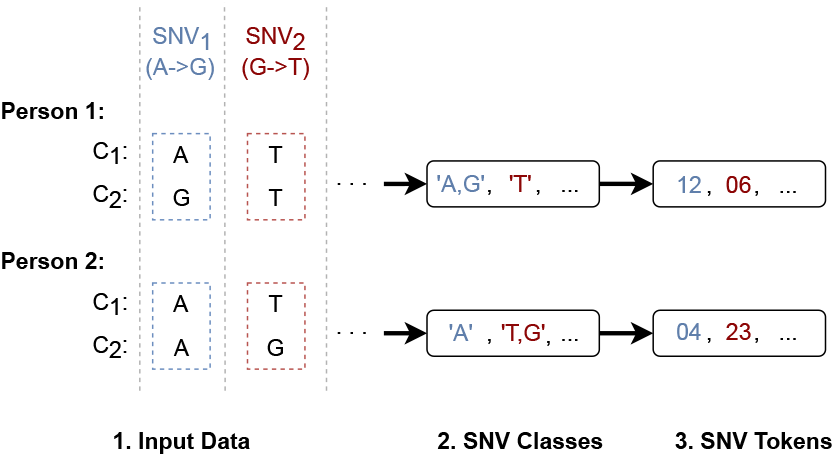
Pre-processing of SNV sequences. Given an input set of SNVs for each person, the combination of SNVs and zygosity is mapped to one of 32 possible cases. Each case is encoded as an integer, and each person’s sequence is encoded as a sequence of these integers.

We separate each SNV into two components, its position and its major/minor alleles. SNV positions are preserved as integer positions within the genome sequence. We mapped different variants and their zygosity to 32 possible combinations as follows. We take the SNV: [*major*]*/*[*minor*] and combine the major and minor alleles into a single word depending on the zygosity, encoding two major, two minor, or one major/one minor allele as follows:

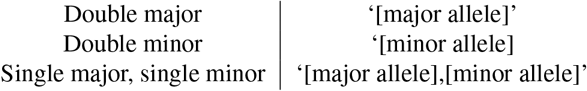

We used insertion (**ins**) and deletion (**del**) tokens to represent any form of insertion and deletion, with respect to the reference genome, in the given SNV. We follow (Cahyawijaya et al., 2022) in shortening long sequences with an **‘I’** token, representing all nucleotides after the first. Doing so allows us to remove the large number of unique long nucleotide sequences from our mapping. A sequence of more than one nucleotide is then encoded as **‘AI’**,**’CI’**,**’GI’** or **‘TI’**, depending on the nucleotide in the first position. In any individual, approximately 2% of SNVs are unknown. That is, we do not know whether either chromosome possesses this variant. We record these as the **‘nan’** token. Finally, we mapped the resulting 32 possible combinations to 32 integers. For example, if we have an SNV that has **‘G, TGAA’** major and minor values, we encode this as **‘G, ins’**, which maps to 21. For the complete list of the 32 combinations, see Table 1.

We classify each individual as either having or not having gout following the method in (Cadzow et al., 2017), with the exception being that classification is now based on self-report or urate-lowering therapy (ULT) only, defined as an individual self-reporting use of allopurinol, probenecid, or colchicine, unless diagnosed with leukemia or lymphoma.

We consider SNVs from the genotyped data only (unimputed), with a minimum allele frequency of 10^−4^, excluding values with a Hardy-Weinberg equilibrium exact test p-value threshold of 10^−6^. Among these, we use plink1.9 (Purcell et al., 2007) to find measured or imputed SNVs associated with urate via linear regression, with a p-value < 10^−1^. The resulting data set contains ≈ 66, 000 SNVs. Since the majority of UKBB participants do not have gout, we randomly sub-sample the non-gout cases until we have ≈ 9, 000 of each. 25% of these are reserved for the test set, with remaining 70% used for training, and 5% saved for later verification. As well as genotype information, we include age, sex, and BMI for each individual.

#### 2.1. Transformer Network Architecture

We embed each SNV as a one-hot encoding of the token (Table 1), concatenated with a sine/cosine positional encoding. This is then passed into a Transformer encoder followed by a binary output block for gout/non-gout classification. Self-attention blocks within the encoder use the Linformer architecture (Wang et al., 2020).

The output takes the classification token (first token of the encoder output), reduces it to two cases with a fully connected layer, and classifies with binary Softmax. Our architecture is summarised in Figure 2.

**Figure 2:**
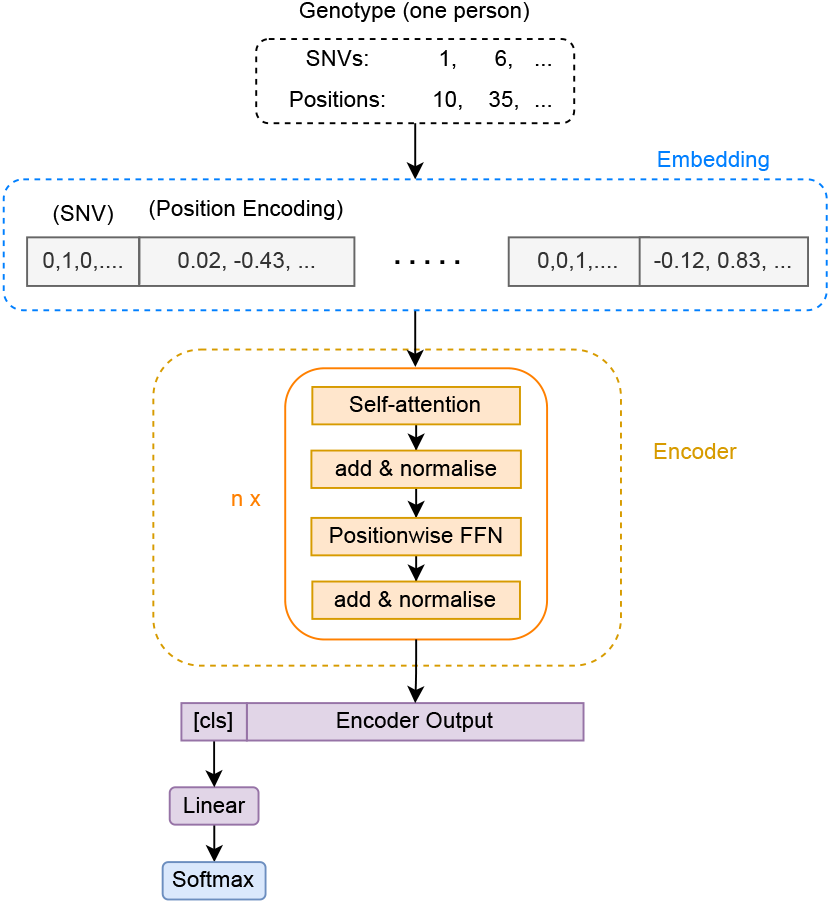
Architecture of SNVformer. Input (genotype) is shown at the top, followed by the embedding, network’s layers, and output at the bottom.

We used a 96 element embedding (where the input uses 32 elements for the token and 64 for the position), 4 attention heads, 6 layers, and a Linformer k-value of 96. We trained using the AMSGrad variant of the AdamW algorithm (Reddi et al., 2019) for 60 epochs with a learning rate of 10^−7^ using batches of size 10 on one 48GB Nvidia Quadro RTX 8000 GPU.

## 3. Results

We evaluate our network using a test set of 5, 364 individuals, exactly half of whom have gout. For each sample, the network outputs a score in the range [0, 1], which we interpret as a prediction that the individual has gout if it is greater than 0.5, and that they do not have gout if it is less than 0.5. The exact score can be interpreted as a confidence indicator for that prediction, and we see in Figure 3b that as scores deviate from 0.5, the accuracy of the prediction increases significantly. Moreover, we see in Figure 3a that scores less than 0.1 or greater than 0.9, where accuracy is around 80%, are not uncommon. We will build on this to perform an analysis of which features are contributing to the certainty and uncertainty of predictions. Based on the output score of the network, we achieve an area under the receiver operating characteristic curve (AUROC) of 0.83 (Figure 4).

**Figure 3:**
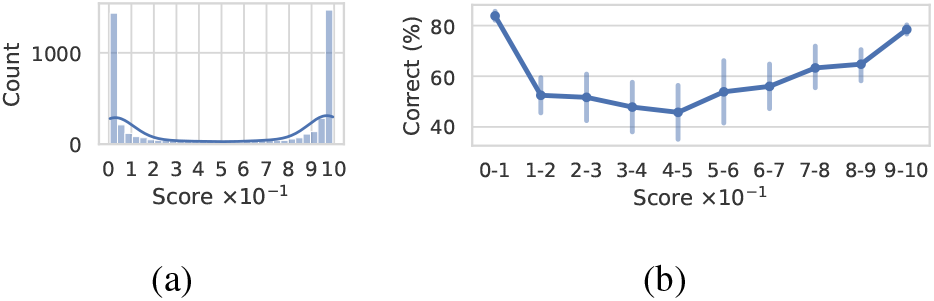
(a) Distribution of output scores produced by our network. Scores can be interpreted as network’s confidence indicator. (b) Comparison between accuracy (vertical) and output score (horizontal) produced by our network. Includes the 95% confidence interval from 1,000 bootstrap samples.

**Figure 4:**
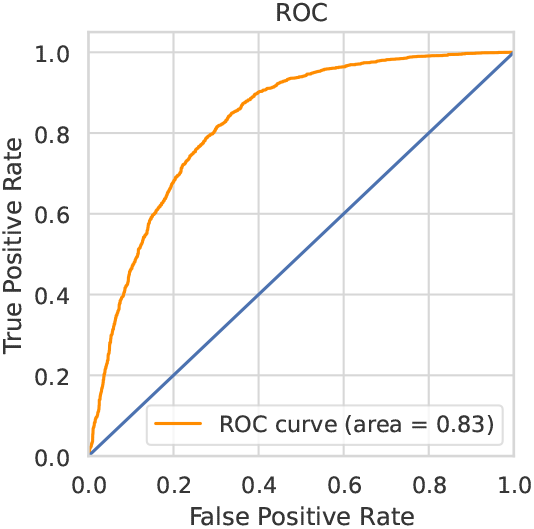
ROC curve when trainied on 66k genotyped SNVs combined with Age, Sex, and BMI.

## 4. Discussion and Work in Progress

Although our preliminary attempt described in this paper has already achieved prediction accuracy similar to the best performing methods (0.84, see below), there are paths to architectural and data-driven improvements.

Other Transformer architectures have been proposed for the problem of learning over long sequences (in NLP). Memorizing Transformers learn to use a fixed-length “short-term” memory to store tokens expected to be useful later (Wu et al., 2022), while Jaegle et al. iterate over read-process-write tasks. The latter approach identifies and queries missing information from the input, processes and accumulates it in a latent space, and uses that latent representation both for prediction and to further query the input. Radically different from Linformer in their approaches to distant dependencies, these architectures may further improve the identification of distant interacting SNVs in long sequences.

Although GWAS analysis provides some explanatory power by identifying SNVs most predictive of gout, it is commonly limited to occurrence or co-occurrence; predictive power could be substantially improved if relevant broader gene-gene (including higher-order) interactions could be identified. A popular method for explaining predictions in Transformers is by analysing the values of the attention matrix, often by visualisation (Vig, 2019), since these reflect the task-dependent *importance* of each token (Wiegreffe & Pinter, 2019). Alternatives include methods to compute and propagate trained attention-based token relevancy scores (Chefer et al., 2021), or to generate higher-level conceptual explanations (Rigotti et al., 2021). We plan to apply such methods to identify which specific SNVs and combinations support phenotype (gout) prediction. We shall also seek to improve the SNV sequence representation for prediction by pretraining embeddings using sequences from the whole UKBB, using large-scale language-modelling techniques, and, based on preliminary experiments, by investigating other encodings for the SNVs themselves.

We intend to compare our method to polygenic risk-scores (Richardson et al., 2019), which for gout and its related traits have achieved prediction accuracy (AUROC) of between 0.64 and 0.91 (Zhang & Lee, 2020; Tin et al., 2019; Sun et al., 2022) depending on whether the model includes additional factors such as demographics and/or clinical factors. For example, Sun et al. have developed a prediction model for clustering the renal underexcretion of urate phenotype using genetic and clinical variables, achieving AU-ROC of 0.91 when 7 clinical measures (age, hypertension, nephrolithiasis, blood glucose, serum urate, urea nitrogen, and creatinine) are added. For gout, prediction accuracy (AUROC) improved from 0.68 to 0.84 with the addition of age and sex (Tin et al., 2019). It will be important to assess how these phenotypic and other variables impact the prediction accuracy of our approach given that certain environmental factors might also modify the effect of genetic risk.

Due to the facility of Transformer models in representing complex language, they have the potential to use a far wider range of supporting data than that used in previous work on gout prediction. In particular, a major focus of our future work will involve enabling our models to access existing well-established evidence of interactions directly from the scientific literature, providing context to the Transformer learner itself. We will apply existing techniques from our labs and others for open domain Transformer-based question-answering against the biomedical literature and therefore extract evidence of genomic interactions, including interactions supported by multi-step protein-protein interaction pathways.

## Acknowledgements

AG acknowledges support from the Royal Society Te Apārangi through a Rutherford Discovery Fellowship (RDF-UOC1702). This work was partially supported by the Ministry of Business, Innovation, and Employment of New Zealand through a Data Science Programmes grant (UOAX1932).

